# Impacts of Spaceflight Experience on Human Brain Structure

**DOI:** 10.1101/2022.02.09.479297

**Authors:** Heather R. McGregor, Kathleen E. Hupfeld, Ofer Pasternak, Nichole E. Beltran, Yiri E. De Dios, Jacob J. Bloomberg, Scott J. Wood, Ajitkumar P. Mulavara, Roy F. Riascos, Patricia A. Reuter-Lorenz, Rachael D. Seidler

## Abstract

Spaceflight induces widespread changes in human brain morphology. It is unclear if these brain changes differ with varying mission durations or one’s history of spaceflight experience (e.g., number of prior missions, time between missions). Here we addressed this issue by quantifying voxelwise post-flight changes in gray matter volume, white matter microstructure, extracellular free water (FW), and ventricular volume in a sample of 28 astronauts. We found that longer missions induced greater ventricular expansion and larger FW displacement at the top of the brain. A greater number of prior missions was associated with white matter microstructure declines in a tract supporting voluntary leg movement. Longer inter-mission intervals were associated with greater ventricle expansion, with compensatory ventricular expansion observed only in those crewmembers with inter-missions intervals of 3 years or longer. Longer missions therefore induce more extensive brain fluid shifts, and the ventricles may require at least 3 years to recover post-flight.

## Introduction

Spaceflight imposes multiple hazards on the human body including increased radiation, microgravity exposure, and social isolation and confinement in a closed environment, among other factors (1). A number of studies have now reported that spaceflight alters human brain morphology (2–15). Spaceflight induces an upward shift of the brain within the skull (13, 16), resulting in cortical crowding and narrowing of the sulci at the top of the brain (4, 16, 17). This is reflected as widespread gray matter volume (GMv) increases at the top of the brain and GMv decreases around the base of the brain (3, 4, 14, 16–18). These post-flight GMv shifts are accompanied by displacement of intracranial fluid (19) including extracellular free water (FW) such as cerebrospinal fluid. Following spaceflight, there are decreases in intracranial fluid volume at the top of the brain and increases around the base of the brain (3, 13, 17, 19, 20). Ventricular expansion also occurs with spaceflight, with reported average volume increases ranging from 11% to 25% (2–4, 6, 9, 11, 12, 16, 17, 19, 21). However, astronauts vary considerably in their current and past spaceflight experience. Mission durations typically range from 2 weeks to 1 year, and some astronauts are novices while others are experienced (with varying numbers of prior flights and inter-mission intervals). It is not known if or how these individual differences in prior flight experience are associated with spaceflight-induced structural brain changes and intracranial fluid shifts. Gaining insight into experience-dependent structural brain changes with spaceflight is crucial with multi-year human missions to Mars on the horizon. Determining whether brain changes continue throughout prolonged microgravity exposure or plateau at some point during flight will help us to better understand the nature and mechanisms of these changes.

Previous studies have leveraged the flight duration differences between short-duration (2 week) space shuttle missions and long-duration (6 months or more) International Space Station (ISS) missions to study the impact of spaceflight duration on brain changes. Six-month missions result in larger GMv shifts than 2-week missions (14, 16), and year-long ISS missions induce even greater GMv increases within sensorimotor cortical regions compared to 6-month missions (17). These findings suggest that longer time in microgravity results in greater cortical crowding at the apex of the brain. Roberts and colleagues also found greater ventricular volume expansion following 6-month flights than 2-week flights (6, 16). Spaceflight results in brain white matter (WM) microstructure changes within tracts subserving vestibular function (18), visual function (11), visuospatial processing (18), and sensorimotor control (4, 18). Longer duration missions result in smaller microstructural changes within the cerebellar WM compared to shorter missions (18). Although it seems counterintuitive that there would be a greater change in this structure for shorter missions, this may reflect an early, adaptive structural change in-flight that gradually returns to baseline over time.

Our group has also reported associations between FW shifts and individual differences in previous spaceflight experience (13). Astronauts who had completed a greater number of previous missions showed FW decreases in the anterior, medial portion of the brain whereas less experienced astronauts showed FW increases in this region (13). This pattern was irrespective of the flight duration and cumulative number of days in space, indicating that the number of gravitational transitions experienced (as opposed to the amount of time spent in space) has an important effect on the brain’s FW distribution (13). It is possible that repeated adaptation to multiple gravitational transitions may affect the gross morphology of the brain. Time between successive missions may also impact spaceflight-induced brain changes. Ventricular enlargement persists after spaceflight, showing only partial recovery in the following 6-12 months (3, 9, 17, 20). We recently reported that astronauts with less recovery time between missions had larger ventricles pre-flight (even after correcting for age effects), and they showed smaller ventricular volume increases with subsequent missions (17). These findings suggest that crewmembers with larger ventricles pre-flight (whether due to older age or prior spaceflight experience) have less available room or compliance for ventricular expansion with spaceflight.

Here we examined whether and how spaceflight-induced brain changes interact with individual differences in current and previous spaceflight experience including: the duration of a mission, whether a crewmember was novice or experienced, their number of previous missions, and time elapsed since a previous mission. We leveraged MRI data from a sample of 28 astronauts who varied along the following flight experience dimensions: mission duration (approx. 2 weeks to 1 year), previous spaceflight experience (0 to 3 previous missions), and time since previous flight (approx. 1 to 9 years). We conducted whole-brain voxelwise analyses assessing associations between these individual differences and changes in GMv, FW fractional volume, FW-corrected WM microstructure, and ventricular volume. In light of previous work, we hypothesized that longer mission durations would induce greater GMv and FW shifts, greater ventricular expansion, and smaller WM microstructure changes. We hypothesized that greater previous flight experience and smaller inter-mission intervals would induce smaller brain structure changes and FW shifts. The latter finding would suggest reduced compliance/elasticity following a prior flight.

## Results

### Spaceflight induces gray matter shifts, free water redistribution, & ventricular enlargement

We assessed group-level pre- to post-flight changes in GMv, ventricular volume, FW fractional volume, and FW-corrected WM diffusion indices. Analyses were adjusted for astronaut age at launch, sex, mission duration, and time elapsed between landing and the post-flight MRI scan. Two-tailed t-test results were thresholded at p < 0.05 with FWE correction.

As shown in the Supplemental Information section, we found statistically significant GMv shifts following spaceflight with apparent GMv increases at the top of the brain and GMv decreases around the base (Figure S1), replicating previous results of our group (14, 17) and others (3, 4, 16). Consistent with previous studies (3, 4, 6, 9, 11, 12, 16, 17, 21), we observed statistically reliable enlargement of the lateral and third ventricles following spaceflight (Figure S2; Table 1). We observed no statistically reliable group level changes in FW-corrected WM diffusion indices from pre- to post-flight. Also replicating our previous work (3, 16–18, 20), spaceflight resulted in widespread decreases in FW fractional volume around the vertex of the brain and FW fractional volume increases around the lower temporal and frontal lobes (Figure S3).

**Table 1.** Pre- to post-flight ventricular volume changes. Beta and p values listed are those corresponding to the variable of interest for each statistical model shown in the leftmost column. Statistically reliable results (p < 0.05) are underlined. Results that additionally survived the Benjamini-Hochberg FDR correction are bolded.

| Model | Ventricle |  |  |  |  |  |  |  |
| --- | --- | --- | --- | --- | --- | --- | --- | --- |
|  | Left Lateral |  | Right Lateral |  | Third |  | Fourth |  |
| | $\beta$ | p | $\beta$ | p | $\beta$ | p | $\beta$ | p |
| Session (pre-flight, post-flight) | 0.603 | <u>0.0004</u> | 0.494 | <u>0.0004</u> | 0.116 | <u>< 0.0001</u> | -0.011 | 0.3434 |
| Current Mission Duration | 0.0039 | <u>0.0180</u> | 0.0036 | <u>0.0062</u> | 0.00056 | <u>0.0004</u> | -0.00012 | 0.3798 |
| Novice vs. Experienced | 0.0073 | 0.9816 | 0.03385 | 0.8976 | 0.0118 | 0.7128 | 0.02389 | 0.3479 |
| Previous Number of Missions | -0.1191 | 0.4392 | -0.0771 | 0.5472 | -0.0101 | 0.5185 | -0.0206 | 0.0899 |
| Years Since Previous Mission End | 0.2367 | <u>0.0481</u> | 0.21517 | <u>0.0314</u> | 0.0305 | <u>0.0081</u> | -0.0120 | <u>0.0488</u> |

### Longer missions induce greater fluid shifts

We tested for pre- to post-flight changes in GMv, ventricular volume, FW fractional volume, or WM diffusion indices that scaled with the current mission duration. Analyses were adjusted for astronaut age, sex, and time elapsed between landing and the post-flight MRI scan. Two-tailed t test results were thresholded at p < 0.05 with FWE correction.

We found no statistically reliable associations between current mission duration and pre- to post-flight GMv shifts.

As shown in Table 1, mission duration was associated with pre- to post-flight increases in left lateral ventricle volume (β = 0.0039, p = 0.018), right lateral ventricle volume (β = 0.0037, p = 0.006), and third ventricle volume (β = 0.00056, p = 0.0004) though only the results for the right lateral and third ventricle survived FDR correction using the Benjamini-Hochberg method (see methods) (22). As shown in Figure 1, 2-week-long missions resulted in smaller increases (or in some instances decreases) in right lateral and third ventricle volume compared to missions lasting 6 months or longer.

**Figure 1.**
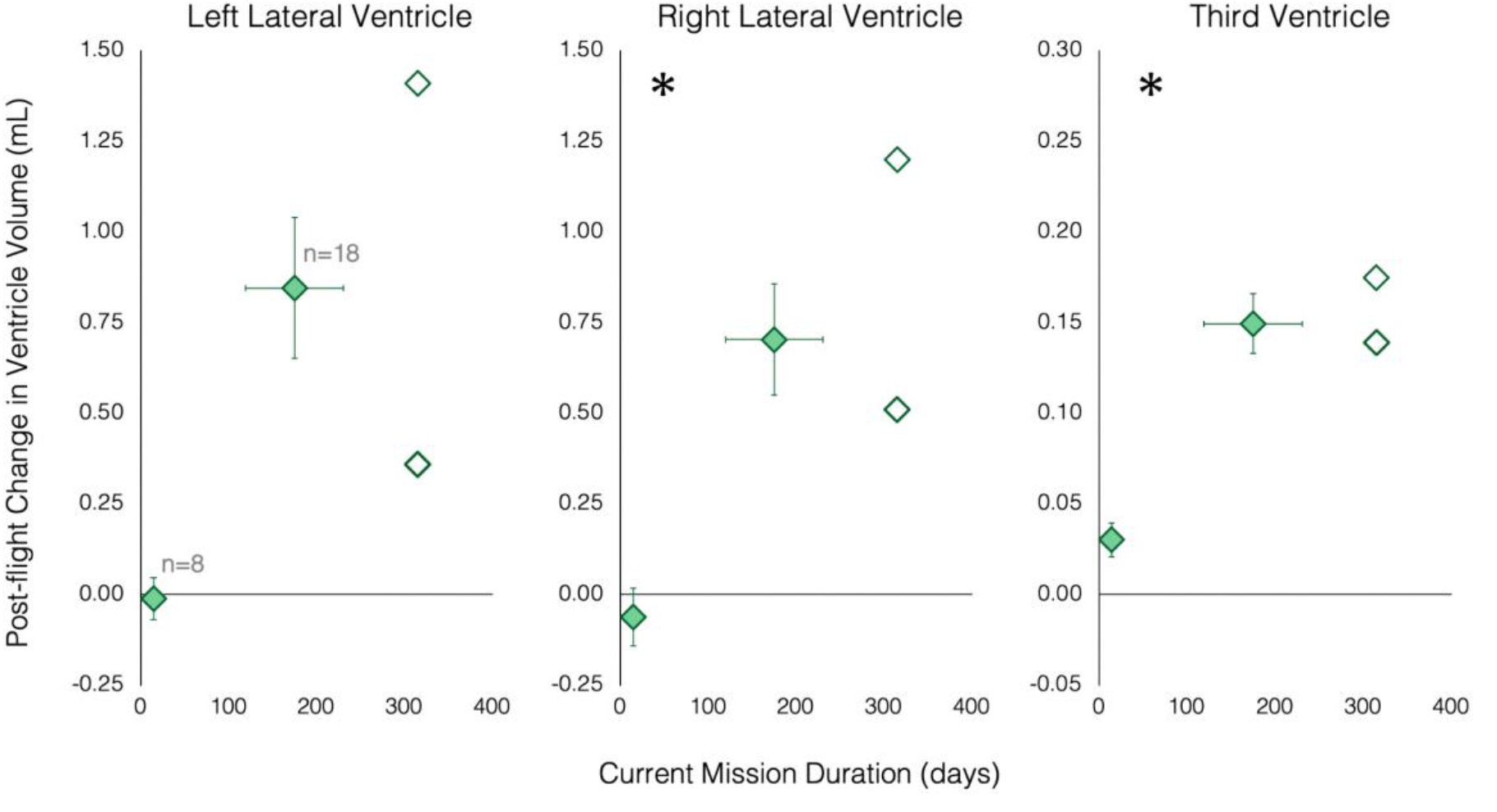
Pre- to post-flight ventricular volume changes associated with mission duration. Filled diamonds represent average values of subgroups of astronauts who completed 2-week-long (n-8) or 6-month-long (n=18) missions. Subgroup sample sizes are indicated in gray on the leftmost plot. Open diamonds represent values for two individual astronauts who completed 1-year-long missions. Asterisks indicate results that survived the Benjamini-Hochberg FDR correction. Error bars represent standard deviation. The two astronauts who completed 1-year missions provided consent for presentation of their individual data.

We also observed an association between mission duration and FW fractional volume changes in clusters located primarily within the precentral, central, and postcentral sulci at the apex of the brain (Figure 2). While 2-week-long missions resulted in FW increases or small decreases within these clusters, missions lasting 6 months or longer resulted in FW fractional volume decreases within these clusters in all but 1 crewmember (who had completed a 6-month-long mission).

**Figure 2.**
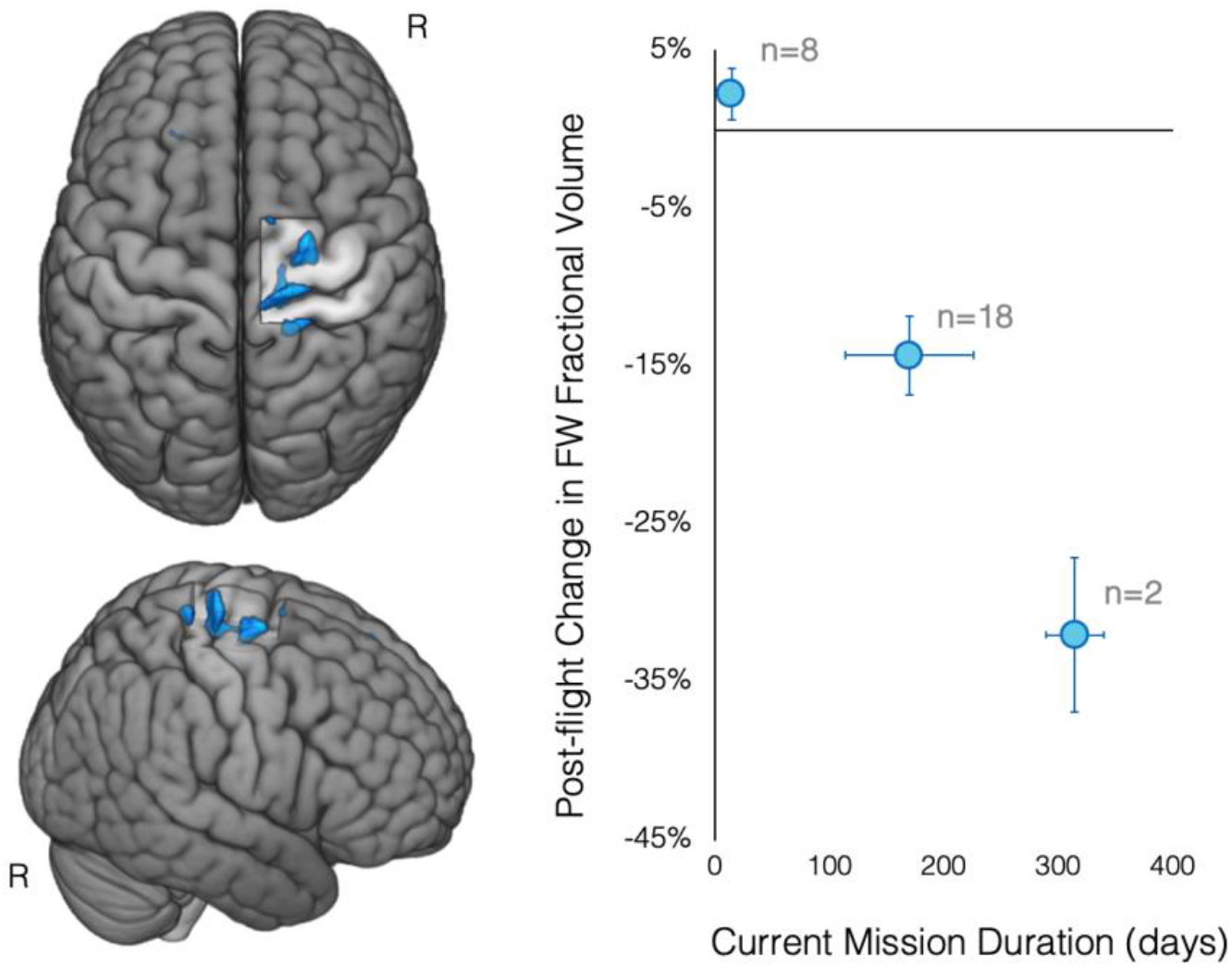
Pre- to post-flight FW fractional volume changes associated with mission duration. Clusters within the right precentral, central, and postcentral sulci are overlaid on a rendered MNI standard space template (left). The scatterplot shows that longer mission durations were associated with greater post-flight FW volume decreases within these clusters. Results are FWE corrected at p < 0.05, two-tailed. Subgroup sample sizes are indicated in gray on the plot. R indicates the right hemisphere. FW, free water.

There were no statistically reliable associations between mission duration and post-flight WM microstructure changes.

### No differences in structural brain changes between novice and experienced astronauts

We tested for differences in post-flight GMv, ventricular volume, FW fractional volume, or WM diffusion index changes between novice and experienced flyers. We categorized astronauts based on whether or not they had any previous spaceflight experience. “Novice” astronauts were those who had no previous spaceflight experience whereas “experienced” flyers were those who had previously completed one or more previous missions. Analyses were adjusted for astronaut age, sex, mission duration, and time elapsed between landing and the post-flight MRI scan. Two-tailed t-test results were thresholded at p < 0.05 with FWE correction.

Whole-brain analyses revealed no statistically reliable associations between post-flight changes in GMv, ventricular volume (see Table 1), FW fractional volume, or WM microstructure (FAt, RDt, ADt) and whether a crewmember was a novice or experienced flyer.

### Greater prior flight history associated with decreases in FW and WM following spaceflight

We then assessed whether the extent of previous spaceflight experience was associated with spaceflight-induced brain changes. We examined pre- to post-flight changes in GMv, ventricular volume, FW fractional volume, or WM diffusion indices that were associated with the number of prior missions completed. Analyses were adjusted for astronaut age, sex, and time elapsed between landing and the post-flight MRI scan. Two-tailed t test results were thresholded at p < 0.05 with FWE correction.

There were no statistically reliable associations between the number of previous missions completed and post-flight GMv shifts or ventricular volume changes.

Voxelwise FW analyses revealed a cluster at the edge of the right lateral ventricle in which post-flight changes in FW fractional volume were associated with the number of previous missions crewmembers had completed. As shown in Figure 3, novice astronauts exhibited FW fractional volume increases within this region following spaceflight. In contrast, crewmembers who had completed multiple previous missions tended to show FW decreases within this region. One crewmember who had completed 2 previous missions was a negative outlier, showing FW decreases more than 5 SD outside of the subgroup mean; this crewmember has been omitted from the plot shown in Figure 3.

**Figure 3.**
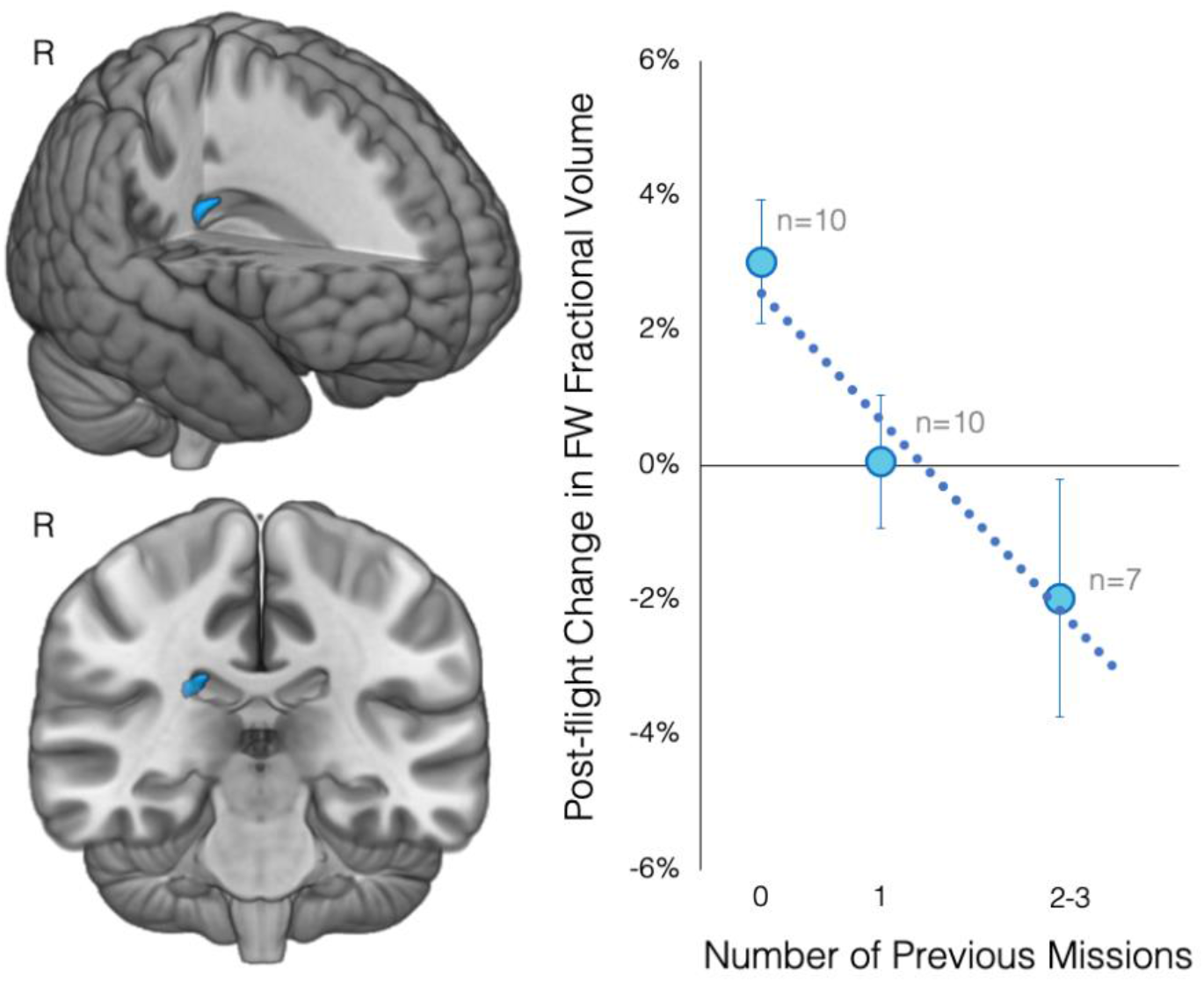
Pre- to post-flight FW volume changes associated with previous number of missions. Cluster located at the outer wall of the right lateral ventricle overlaid on a rendered MNI standard space template (left). The scatterplot shows that having completed a greater number of previous missions was associated with greater post-flight FW volume decreases within this cluster. The dotted line indicates the linear fit of the individual data points. Circular markers indicate the average of subgroups, the size of each subsample is indicated in gray. The 2 and 3 previous missions subgroups have been combined for crewmember privacy. Error bars represent standard deviation. Results are FWE corrected at p < 0.05, two-tailed. R indicates the right hemisphere. FW, free water.

Our analyses also identified that the previous number of missions completed was associated with one measure of WM microstructure change following spaceflight (Figure 4). Specifically, astronauts with greater previous spaceflight experience showed decreases in FW-corrected axial diffusivity (ADt, indicating the magnitude of diffusion along WM fibers) in a small cluster within the right posterior corona radiata. Novice and less experienced astronauts tended to show ADt increases within this region.

**Figure 4.**
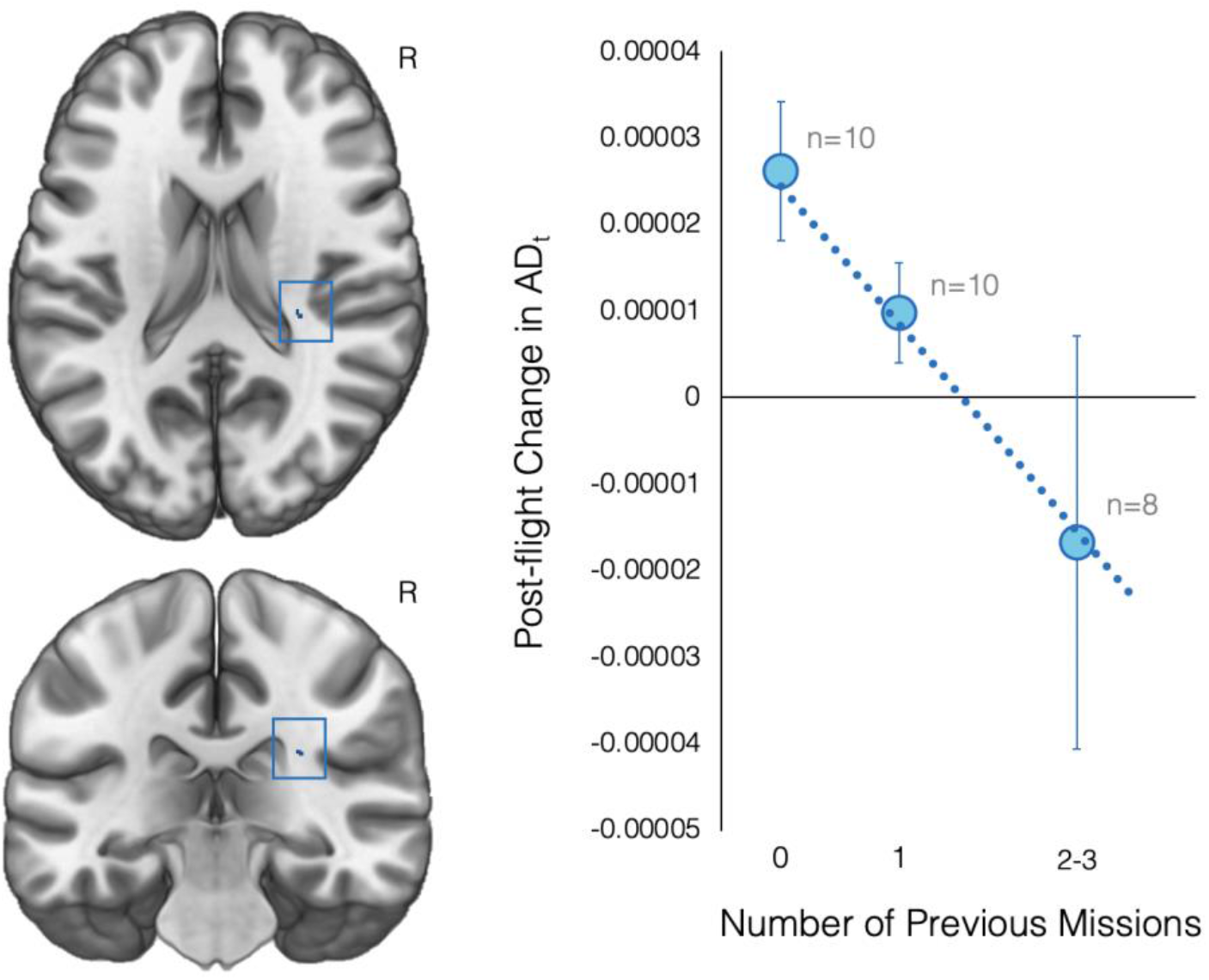
Pre- to post-flight AD_t_ changes associated with previous number of missions. Cluster within the right posterior corona radiata overlaid on a rendered MNI standard space template. The plot shows that having completed a greater number of previous missions was associated with greater post-flight AD_t_ decreases within this cluster. The dotted line indicates the linear fit of the individual data points. Circular markers indicate the average of subgroups, the size of each subsample is indicated in gray. The 2 and 3 previous missions subgroups have been combined for crewmember privacy. Error bars represent standard deviation. Results are FWE corrected at p < 0.05, two-tailed. R indicates the right hemisphere. ADt, free water-corrected axial diffusivity.

Analyses yielded no statistically reliable associations between the number of previous missions completed and the other FW-corrected WM diffusion indices examined.

### Greater ventricular expansion for astronauts with longer inter-mission intervals

We tested whether post-flight changes in GMv, ventricular volume, FW fractional volume, or WM diffusion indices scaled with the time interval between successive missions. Analyses were adjusted for astronaut age, sex, and time elapsed between landing and the post-flight MRI scan. Two-tailed t test results were thresholded at p < 0.05 with FWE correction.

We found no statistically reliable associations between the time elapsed since the previous mission and post-flight GMv shifts.

Among the experienced astronauts, the number of years elapsed since the previous mission was significantly associated with post-flight volume changes for all four ventricles (see Figure 5). Longer time between successive missions was associated with greater increases in left lateral (β = 0.23668, p = 0.0481), right lateral (β = 0.21517, p = 0.0314), and third ventricle (β = 0.0305, p =0.0081) volumes following spaceflight. The fourth ventricle showed the opposite pattern with longer intermission delays being associated with greater volumetric decreases following spaceflight (β = −0.0120, p = 0.0488). However, only the association between third ventricle volume changes and years since the previous mission survived FDR correction using the Benjamini-Hochberg correction (22). For experienced crewmembers, shorter intermission intervals (typically < 3 years) were associated with smaller increases (or in one astronaut a decrease) in the volume of the third ventricle. Longer intermission intervals were associated with larger expansion of the third ventricle following spaceflight. These results are also presented in Table 1 and Figure 5.

**Figure 5.**
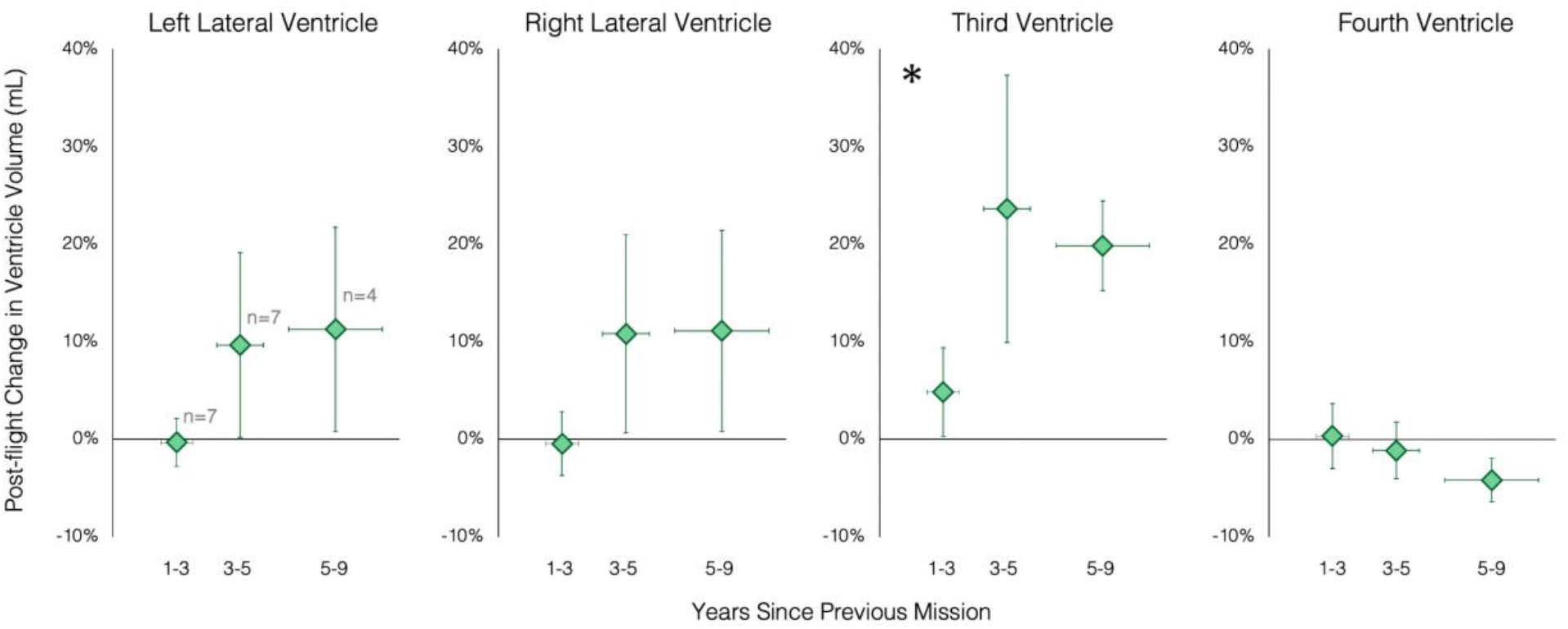
Pre- to post-flight ventricle volume changes associated with inter-mission intervals. Pre- to post-flight ventricle volume changes of the third ventricle associated with the number of years since the previous mission’s end. Crewmembers have been subgrouped based on inter-mission intervals for crewmember privacy. The x axis shows exclusive ranges (i.e., encompassing intermission intervals up to, but excluding the upper bound). Markers indicate subgroup averages with subgroup sample sizes indicated in gray in the leftmost plot. Error bars indicate standard deviation. Asterisks indicate results that survived the Benjamini-Hochberg FDR correction.

There were no statistically reliable associations between time since the previous mission and changes in FW volume or FW-corrected WM diffusion indices.

## Discussion

Here we showed that longer duration spaceflight resulted in greater ventricular enlargement and larger reductions in FW from within the sulci at the top of the brain. Crewmembers with greater prior spaceflight experience showed white matter microstructure declines within tracts subserving voluntary leg movement. Longer recovery time between subsequent missions was associated with greater post-flight ventricular expansion. This work suggests that longer missions, multiple flights, and shorter inter-mission recovery time induce greater intracranial fluid shifts and focal WM decline.

### Associations with Mission Duration

Longer duration missions were associated with greater enlargement of the right lateral ventricle and third ventricle following spaceflight. Our data show that the right lateral ventricle may continue to expand during missions lasting longer than 6-month missions whereas expansion of the third ventricle begins to taper off during 6-month-long missions. Lateral ventricle volumetric changes were heavily influenced by one of the astronauts on a 1-year mission who exhibited large pre-flight ventricles and showed small ventricular volume changes following the current year-long mission. This crewmember’s ventricles may not have had room or sufficient elasticity to expand to a greater extent during spaceflight (17). Our 1-year mission duration data were based on only 2 astronauts both of whom had previous spaceflight experience. As such, this finding should be interpreted with caution since carryover effects between missions remain poorly understood.

We also observed that longer duration missions induced larger FW fractional volume decreases within the sulci neighboring the pre- and postcentral gyri. This is consistent with a previous qualitative report of sulcal narrowing at the top of the brain following long-duration spaceflight (16). This FW volume change likely reflects expulsion of cerebrospinal fluid from between the folds at the top of the brain. These FW volume shifts showed no indication of plateauing across the mission durations examined, suggesting that missions lasting longer than 1 year will result in even more extensive narrowing of cerebrospinal fluid (CSF) spaces at the top of the brain. Thus, there could be a risk of even greater impedance of CSF flow and/or resorption at the top of the brain during multi-year missions. Examination of structural brain changes of additional crewmembers yearlong missions is required to characterize CSF displacement at the top of the brain and determine at what mission duration these FW volume changes begin to plateau.

### Associations with Previous Flight Experience

Novice crewmembers showed FW fractional volume increases at the edge of the right lateral ventricle following spaceflight while those with experience of 2 or more previous missions showed FW fractional volume decreases within this region. Using a subset of the prospective astronaut data, we previously showed that crewmembers who had completed more previous missions showed smaller enlargement of the right lateral ventricle (17). This may be due to cumulating carryover effects from previous flights. In-flight ventricular expansion and slow recovery following spaceflight (2, 9, 17) may reduce the compliance of the ventricles thus reducing their efficacy as an overflow zone for fluid shifts and reduced CSF resorption during missions (2, 7, 21). This would be especially the case if the brain does not fully recover between flights and crewmembers begin a subsequent flight with enlarged ventricles (17), which may have been the case for one of the 1-year crewmembers noted above.

Additionally, we found that previous spaceflight experience was associated with greater decreases in WM microstructure within the posterior portion of the corona radiata. The corona radiata is a major WM tract projecting to and from the cerebral cortex. ADt reflects the magnitude of diffusion along WM fibers. AD is a sensitive measure of axon degeneration (23, 24), however, this assumes that the WM fibers are coherently aligned and no tissue realignment has occurred (24). Given the complex nature of brain tissue shifts, fluid redistribution, and ventricular expansion occurring during spaceflight, we interpret this particular finding with caution. Repeated in-flight expansion and post-flight contraction of the lateral ventricles may exert mechanical strain on this neighboring WM tract, thus causing the observed WM microstructure changes. However, the posterior portion of the corona radiata houses descending cortical motor fibers supporting voluntary leg movement (25, 26). This fact raises the possibility that the observed WM microstructure changes occurred as a result of repeated periods of prolonged leg disuse in-flight. In microgravity, the body is unloaded (i.e., “weightlessness”) and the large muscles in the legs no longer need to continuously contract to support one’s body weight against gravity. In-flight reductions in motor outflow to the legs may underlie the observed ADt increases. Indeed, clinical MRI studies have similarly shown reductions in WM integrity within cortical tracts in patients experiencing limb impairment following spinal cord injury (27). Future studies comparing in-flight leg usage and post-flight WM changes are required to further test this hypothesis. Interestingly, post-flight decreases in ADt within the posterior corona radiata appear to plateau, with crewmembers who had completed 2 or 3 prior missions exhibiting comparable ADt decreases within this WM tract.

It is worth noting that prior flight experience may increase with crewmember age, however all analyses adjusted for individual differences in crewmember age (in addition to sex, current mission duration, etc.); therefore this finding is not attributable to age differences. Our findings suggest that repeated adaptation to changing gravity conditions experienced across multiple previous flights alters WM microstructure and FW fractional volume.

We found no statistically reliable differences in spaceflight-induced brain changes between astronauts when categorizing them on a binary basis as novice or experienced. Instead, differential brain changes emerged when we factored in the number of prior flights the experienced astronauts had completed. This finding suggests that the brain is impacted by the cumulative effects across multiple flights and perhaps separate bouts of adaptation to microgravity and the spaceflight environment.

### Associations with Inter-mission Intervals

Among the experienced astronauts, crewmembers who had less than 3 years of time to recover following their previous mission showed little to no enlargement of the lateral and third ventricles following the current mission. In contrast, those crewmembers who had 3 years or longer to recover following their previous mission showed ventricular expansion following the current mission. Ventricular expansion resolves slowly post-flight (2, 17), with crewmembers showing an average of ~55-64% recovery towards pre-flight levels 6-7 months following a 6-month ISS mission (2, 17). Incomplete ventricular recovery between flights may reduce the brain’s compliance and negatively impact compensatory mechanisms. That is, ventricular expansion during spaceflight may allow the brain to manage the headward fluid shifts that occur with microgravity. Qualitatively, in the current study, most of the crewmembers with inter-mission intervals of 3 years or longer exhibited post-flight lateral and third ventricle expansion (see Figure 5). This suggests that crewmembers may require a 3-year interval between successive missions for post-flight ventricular recovery and regaining compensatory capacity. Using a subset of the prospective data, our group previously reported that less time between successive missions was associated with larger pre-flight lateral ventricles (an effect that was not attributable to age) (17). Analysis of pre-flight ventricle volumes and intermission intervals using the larger dataset in the current study did not replicate this effect, resulting in mixed evidence. We previously reported that less time between successive missions was associated with smaller ventricular volume increases with flight (17). We observed a similar effect here, suggesting incomplete recovery between successive flights. Forthcoming multi-year longitudinal studies will shed more light on post-flight ventricular recovery.

A limitation of this work includes differing MRI scan parameters for a subset of diffusion-weighted MRI images for the retrospective subjects (see methods), with some scan parameters differing between pre- and post-flight scans. Similar to previous studies (2, 3, 6–8, 13–15), post-flight MRI scans occurred an average of 6.5 days following landing (range: 1-20 days). It is possible that some spaceflight-induced brain changes recovered prior to the post-flight brain scan or were impacted by readaptation to Earth’s gravity. Although we adjusted for the time delay between landing and the post-flight MRI scan session in all analyses, this time delay was longer for astronauts in the retrospective dataset. Prospective studies could also apply more advanced dMRI acquisition approaches such as multi-shell and multi-band which would make the model estimations more accurate and will shorten the length of the acquisition.

Here we reported how current and previous spaceflight experience relates to spaceflight-induced brain structural changes. Longer duration missions induced greater ventricular expansion and larger reductions in FW from within the sulci at the top of the brain. A greater number of prior missions was associated with WM microstructure decreases within a WM tract supporting voluntary leg movement. Longer inter-mission intervals were associated with greater post-flight ventricular expansion. This work suggests that longer missions, multiple flights, and shorter inter-mission recovery time induce greater intracranial fluid shifts and focal WM decline. With human spaceflight becoming more frequent and lengthy, these findings provide important insight into how spaceflight experience, both current and previous, impacts the brain.

## Materials and Methods

### Participants

T1-weighted and diffusion-weighted MRI (dMRI) scans collected from a total of 28 astronauts were included in this study. Data from 15 of the astronauts were collected as part of a prospective study conducted between 2014 and 2020 (28). Astronauts in the prospective group completed a long-duration mission to the ISS lasting approximately 6 (n=13) or 12 months (n=2). Data from the remaining 13 astronauts were obtained from the NASA Lifetime Surveillance of Astronaut Health Repository. We obtained MRI scans from 26 crewmembers from this database. Data from 13 of these astronauts were excluded due to missing pre-flight T1 or dMRI scans (n=10), incomplete brain coverage (n=1), insufficient number of diffusion-weighted volumes acquired (n=1), or participation in our prospective study (n=1). Astronauts in the retrospective group completed a short-duration mission lasting approximately 2 weeks (n=8) or a long-duration ISS mission lasting approximately 6 months (n=5). Crewmember demographic information is presented in Table 2.

**Table 2.**
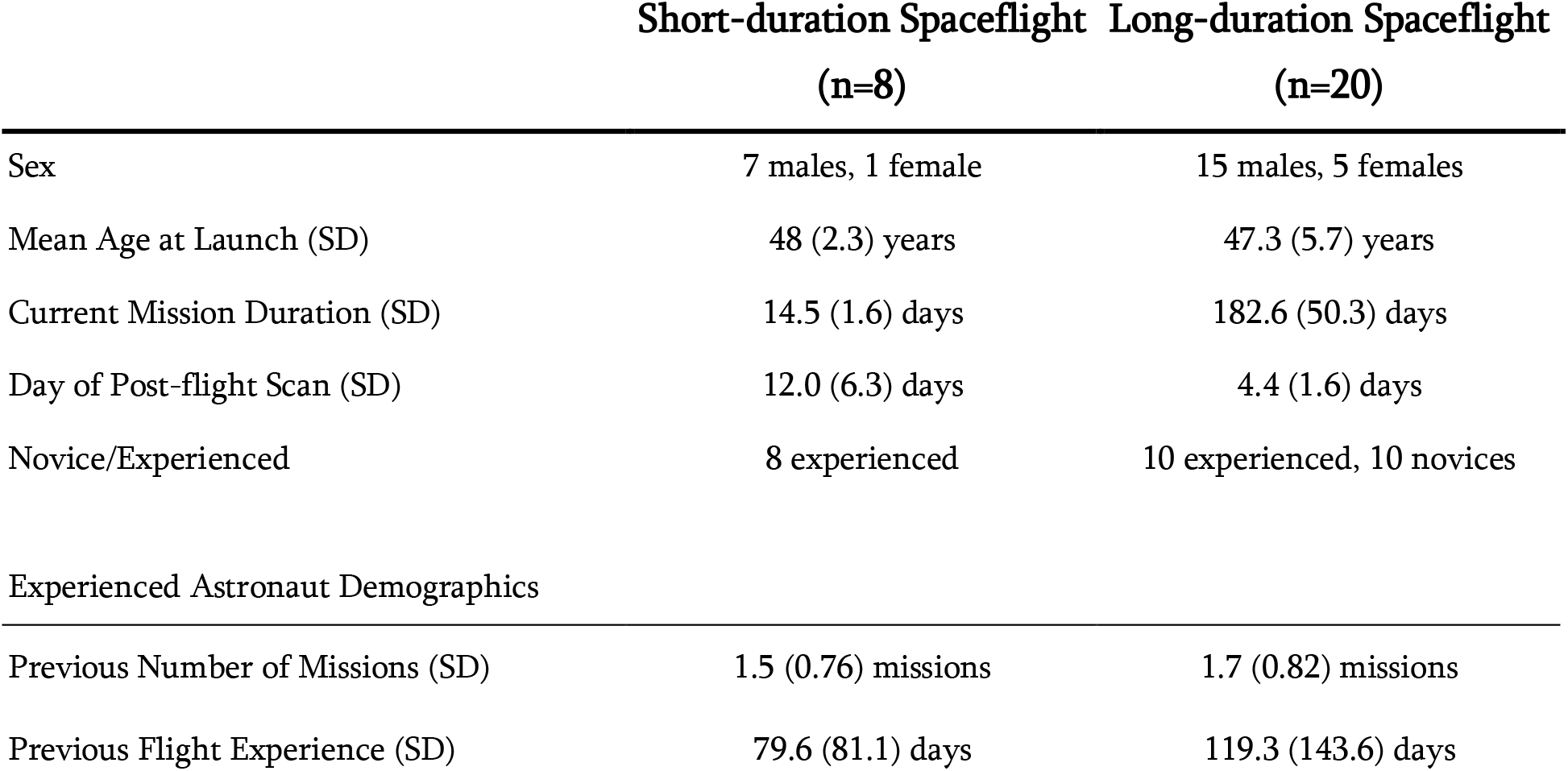
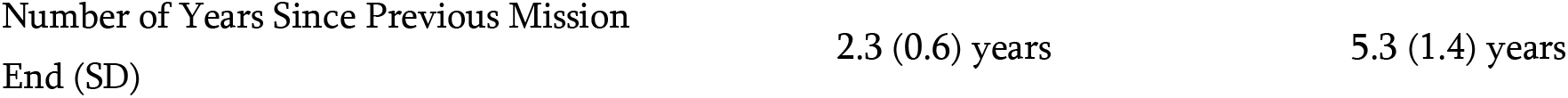
Astronaut Demographics. SD, standard deviation. Experienced refers to having completed at least 1 previous spaceflight mission prior to enrollment in the current study.

|  | Short-duration Spaceflight<br>(n=8) | Long-duration Spaceflight<br>(n=20) |
| --- | --- | --- |
| Sex | 7 males, 1 female | 15 males, 5 females |
| Mean Age at Launch (SD) | 48 (2.3) years | 47.3 (5.7) years |
| Current Mission Duration (SD) | 14.5 (1.6) days | 182.6 (50.3) days |
| Day of Post-flight Scan (SD) | 12.0 (6.3) days | 4.4 (1.6) days |
| Novice/Experienced | 8 experienced | 10 experienced, 10 novices |
| Experienced Astronaut Demographics |  |  |
| Previous Number of Missions (SD) | 1.5 (0.76) missions | 1.7 (0.82) missions |
| Previous Flight Experience (SD) | 79.6 (81.1) days | 119.3 (143.6) days |

This study was approved by the institutional review boards at the University of Michigan, University of Florida, and the NASA Johnson Space Center. Crewmembers provided written informed consent prior to participating in the prospective study.

### MRI Acquisition

T1-weighted anatomical scans and diffusion-weighted MRI (dMRI) scans were acquired pre- and post-flight. All neuroimaging data were acquired using the same 3T Siemens Magnetom Verio MRI scanner located at University of Texas Medical Branch at Victory Lake in Houston, TX.

T1-weighted anatomical images for all astronauts were collected using a magnetization-prepared rapid gradient-echo (MPRAGE) sequence with the following parameters: TR = 1900 ms, TE = 2.32 ms, flip angle = 9°, FOV = 250 x 250 mm, 176 sagittal slices of 0.9 mm thickness, matrix = 512 x 512, voxel size = 0.488 x 0.488 x 0.9 mm.

dMRI scans were acquired using a 2D single-shot spin-echo prepared echo-planar imaging sequence. Prospective dMRI scans were acquired using the following parameters: TR = 11300 ms, TE = 95 ms, flip angle = 90°, FOV = 250 × 250 mm, matrix size = 128 × 128, 40 axial slices of 2 mm thickness (no gap), voxel size = 1.95 x 1.95 x 2 mm. Thirty non-collinear gradient directions with diffusion weighting of b = 1000 s/mm^2^ were sampled twice. A volume with no diffusion weighting (b = 0 s/mm^2^) was acquired at the start of each sampling stream.

Retrospective dMRI scans were acquired using the following parameters: TR = 5800 ms, TE = 95 ms, flip angle = 90°, FOV = 250 × 250 mm, matrix size = 128 × 128, 40 axial slices of 3.9 mm thickness (no gap), voxel size = 1.95 x 1.95 x 3.9 mm. Twenty non-collinear gradient directions with diffusion weighting of b = 1000 s/mm^2^ were sampled three times. At the beginning of each sampling stream, a volume with no diffusion weighting (b = 0 s/mm^2^) was acquired. Voxel dimensions were altered for 9 retrospective crewmembers resulting in a voxel size of 1.8 x 1.8 x 3.9 mm for 9 preflight and 1 post-flight scans. Repetition times were also altered for 4 retrospective crewmembers as follows: 4 pre-flight scans (TR = 5846, 5500, 5900, 5600 ms) and 1 post-flight scan (TR = 5302 ms).

### T1-weighted Image Preprocessing

All T1-weighted images underwent identical preprocessing using the Computational Anatomy Toolbox (29) (CAT12.6 v.1450) for Statistical Parametric Mapping (30) version 12 (SPM12 v.7219) implemented using MATLAB R2016a, version 9.0.

After visual inspection, native space T1 images were skull stripped using CAT12. We additionally segmented each native space T1 image into gray matter (GM), white matter (WM), and cerebrospinal fluid (CSF) segments. GM segments were used in subsequent analyses of gray matter volume (GMv).

CAT12 automatically estimated the volumes, in mL, of the left lateral, right lateral, third, and fourth ventricles using the Neuromorphometrics volume-based atlas map included in SPM12. Ventricular volume analyses were performed in native space. CAT12 additionally provided estimates of each astronaut’s total intracranial volume. Pre-flight total intracranial volumes were included in GMv and ventricular volume analyses detailed below.

### dMRI Preprocessing

All dMRI scans underwent identical preprocessing and analyses using FMRIB Software Library (FSL) version 6.0.1 (31), and a custom FW algorithm (32) implemented in MATLAB R2018b. Raw dMRI scans were visually inspected for scan artifacts and excessive head movement. We used FSL’s preprocessing tool, eddy, to correct for eddy current distortions and inter-volume head movement. Diffusion-weighted volumes (b=1000 s/mm^2^) were registered to the average of the b=0 volumes in the run. Rotations applied to each volume during motion correction were also applied to corresponding diffusion gradient directions. A volume was deemed an outlier if the root mean square voxel displacement was greater than 1 mm relative to the previous volume. Outlier volumes and those with artifacts were removed from the eddy corrected image and from b-value and b-vector matrices. Preprocessed dMRI data were then skull stripped using FSL’s brain extraction tool, bet.

FW maps and FW-corrected diffusion indices were computed by fitting a bi-tensor model at each voxel (32). This model consists of FW and tissue compartments. The FW component models water molecules that are free to diffuse as isotropic diffusion with a diffusion coefficient of water at body temperature (3 x 10^−3^ mm^2^/s). The tissue compartment, modeled using a diffusion tensor, estimates the diffusivity of water molecules within tissue. These components yield FW maps and FW-corrected diffusion indices, respectively. FW maps reflect the fractional volume of the FW compartment within each voxel; these values range from 0 to 1, where 1 indicates that a voxel is filled entirely by freely-diffusing water molecules. The tissue compartment yields the following diffusion index maps: FW-corrected fractional anisotropy (FA_t_), FW-corrected axial diffusivity (AD_t_), and FW-corrected radial diffusivity (RD_t_) that are calculated from the diffusion tensor.

### Image Normalization

As in our previous work (13, 14, 17), we used a multi-step process to improve within-subject registration of our longitudinal images (33). Registration was performed using Advanced normalization tools (ANTs) version 1.9.17 (34, 35). As described below, this multi-step approach involves registering a subject’s native space images to standard space via subject-specific templates.

We first generated subject-specific T1 templates. Subject-specific T1 templates were created by averaging the subject’s native space skull-stripped pre-flight and post-flight T1 images using antsMultivariateTemplateConstruction.sh. In this way, the template was not biased towards either time point. Each subject-specific T1 template was then warped to a 1 mm-resolution MNI standard space T1 template using rigid, affine, and Symmetric Normalization (SyN) transformations using ANTs. Using the method, we also created subject-specific FA templates using subject’s FA images that were not FW-corrected. Native space FA maps were eroded by 1 voxel to remove noise around the outer edge of the volume. Subject-specific FA templates were normalized to an FA template in MNI space as above.

The above steps generated transformations for registering each native space image (T1 or FA) to a subject-specific template, and from a subject-specific template to MNI space. We concatenated these transformations into flow fields to minimize the number of interpolations performed during normalization. A separate flow field was generated for each native space T1 image and FA map. Since FA, FW, FA_t_, AD_t_, and RD_t_ maps from a given session were derived from the same dMRI image, the same flow field was applied to transform each into MNI standard space. These steps yielded MNI-normalized GM segments, FW maps, and diffusion index maps with a resolution of 1mm^3^.

### GM Modulation

As in our previous work (14), flow fields that were applied to GM segments were additionally used to estimate tissue expansion and shrinkage following spaceflight. Each flow field was inputted to ANTs’ CreateJacobianDeterminantImage.sh function to obtain the Jacobian determinant image. The Jacobian determinant encodes local expansion and shrinkage for each voxel within the image. We then multiplied each MNI-normalized GM segment by its corresponding Jacobian determinant image to produce modulated GM segments in standard space for each subject and session. Modulation preserves the amount of GM present in the untransformed image.

### Masking

Voxelwise FW analyses were performed within a whole-brain mask including the ventricles and CSF around the brain parenchyma. Analyses of WM diffusion indices were confined to the WM. A binary WM mask was created by thresholding all MNI-normalized FA maps (≥ 0.2) to identify WM, and included only those voxels in which WM was present in more than half of the sample (13).

### Smoothing

Modulated GM segments were smoothed using an 8 mm (full width at half maximum) FWHM Gaussian kernel to increase signal-to-noise ratio. MNI-normalized FW maps and FW-corrected diffusion indices were each smoothed using a 5 mm FWHM Gaussian kernel.

### Group Level Analyses

#### Statistical Models

Our first model tested for significant brain changes from pre- to post-flight averaged across all astronauts. We then used 4 models to examine associations between pre- to post-flight brain changes and the following crewmember individual differences: current mission duration, whether crewmembers were experienced or novice, previous number of missions (excluding the current mission), and years since previous mission end (for experienced crewmembers only).

Models adjusted for individual differences in age at the time of launch, sex, current mission duration, and the number of days between landing and the post-flight MRI scan (except for models in which one of these covariates was the predictor of interest). GMv and ventricular volume analyses also adjusted for pre-flight total intracranial volume.

#### Whole-brain Voxelwise Analyses

For each astronaut, we calculated FW difference images reflecting pre- to post-flight changes in FW fractional volume. We then concatenated the astronauts’ FW difference images into a single 4-dimensional image. This procedure was repeated individually for each image type.

We performed one-sample t tests on the GMv, FW, FA_t_, RD_t_, AD_t_ difference images using randomise, FSL’s tool for nonparametric permutation-based inference (36). All tests were performed using 15,000 random permutations with threshold-free cluster enhancement (37). Correction for multiple comparisons was implemented using familywise error (FWE) correction (p < .05, two-tailed).

#### Ventricular Volume Analyses

We analyzed pre- to post-flight changes in ventricular volumes using linear mixed models using restricted maximum likelihood (REML). Analyses were implemented in R version 3.3.3 using the nlme package (38). Our linear mixed models modeled random effects corresponding to subject-specific intercepts and slopes. Time was modeled as a categorical variable (i.e., pre-flight or post-flight session). Since we performed a total of 20 ventricular volume analyses (5 analyses performed for each of our 4 ventricle ROIs), we used the Benjamini-Hochberg procedure to control the FDR at p < 0.05 (22).

## Acknowledgments

The authors wish to thank all the crewmembers who volunteered their time to participate in the study. Thank you to Sara Stroble for facilitating retrospective data access.

