## Supplementary material for "Impacts of Spaceflight Experience on Human Brain Structure": Figure S1, Figure S2, Figure S3

### Supplemental Materials

#### Group average pre- to post-flight brain changes

We replicated previous findings showing average pre- to post-flight changes in GMv and FW fractional volume likely reflecting an upward shift of the brain within the skull, cortical compression, and concomitant fluid shifts. As in previous spaceflight (3, 4, 14, 16–18) and bed rest analog studies (38) , we observed GMv shifts from the base of the brain to the apex. We also observed pre- to post-flight FW shifts from the top to the base of the cerebrum, as previously observed with spaceflight (3, 13, 17, 20) and bed rest (39). We additionally replicated previous findings of post-flight enlargement of the lateral and third ventricles (2–4, 6, 9, 11, 12, 16, 17, 19).

We observed no statistically reliable changes in WM microstructure across the brain from pre- to post-flight at the group level. This is in contrast to past research from our group and others showing that spaceflight induces WM microstructural changes within tracts involved in vestibular function (4, 18), visual function (11), visuospatial processing (18), and sensorimotor control (18).

We assessed pre- to post-flight changes in GMv, ventricular volume, FW fractional volume, and FW-corrected WM indices. Analyses were adjusted for astronaut age, sex, mission duration, and time elapsed between landing and the post-flight MRI scan. Two-tailed t test results were thresholded with a familywise error (FWE) correction value p of 0.05.

Replicating previous results of our group (14, 17) and others (3, 4, 16), we found statistically significant GMv shifts following spaceflight (Figure S1), with apparent GMv increases occurring largely within the interhemispheric fissure and surrounding sulci at the apex of the brain and in the parietal operculum, basal ganglia and thalamus. GMv decreases were observed in the right temporal lobe and bilaterally within occipital regions.

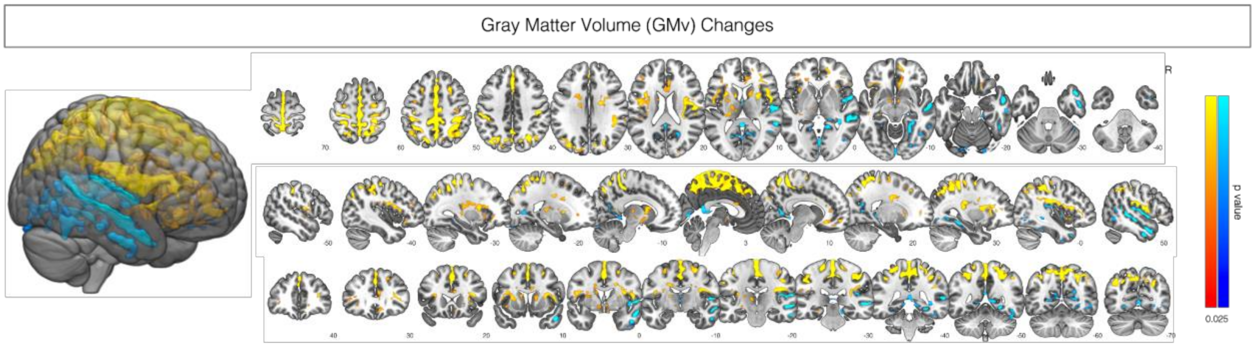


**Figure S1. GMv shifts following spaceflight.** Yellow clusters indicate regions where GMv increased from pre- to post-flight. Blue clusters indicate regions where GMv decreased from pre- to post-flight. Clusters are shown overlaid on a rendered MNI standard space template. Brighter colors indicate smaller p values. Clusters were FWE corrected at $p < 0.05$, two tailed. R indicates the right hemisphere.

Consistent with previous studies (3, 4, 6, 9, 11, 12, 16, 17, 19), we observed statistically reliable enlargement of the lateral and third ventricles following spaceflight (Figure S2). The observed volume increases were between 8 and 16%, which are in range of those previously reported (3, 4, 6, 9, 11, 12, 16, 17, 19). While the group showed post-flight ventricular volume increases on average, it is worth noting that this was not the case for all crewmembers. Of our sample of 28 crewmembers, 6 showed left lateral ventricle volume decreases (all < 4% decreases), 4 showed right lateral ventricle volume decreases (all < 6% decrease), and 1 showed a decrease in third ventricle volume of less than 1%.

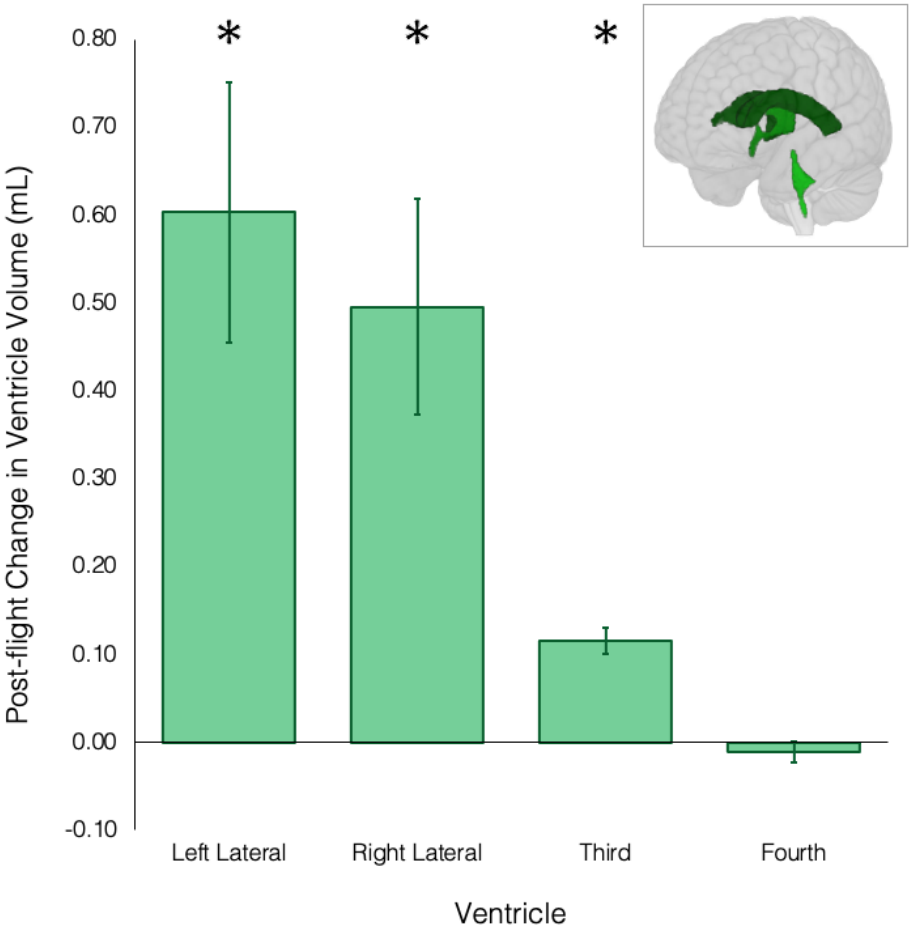


**Figure S2. Ventricular volume changes following spaceflight.** The left and right lateral ventricles exhibited the largest post-flight volume changes, with group average increases of 0.6 mL (8.3%) and 0.5 mL (8.5%), respectively. The third ventricle also showed a statistically reliable post-flight volume increase of approximately 0.1 mL on average (18%). Error bars represent standard deviation. Asterisks indicate results that survived Benjamini-Hochberg FDR correction at $p < 0.05$. The inset at the top right shows the ventricles overlaid on a rendered MNI standard space template. The lateral ventricles are shown in dark green, the third ventricle is shown in medium green, and the fourth ventricle is shown in light green.

Ventricles enlarge over time as part of the normal aging process (40); however, the rate of ventricular expansion during spaceflight exceeds that seen with aging (7, 9, 17). Several groups have reported no changes in global GM and WM volume following spaceflight (2, 6, 12, 14) (though see (9)). This suggests that post-flight ventricular enlargement is not a consequence of brain atrophy with normal aging, but instead arises from CSF changes. While the precise mechanism underlying this phenomenon remains unclear, disruptions in the CSF dynamics have been hypothesized as an underlying cause. Headward fluid shifts in microgravity may impair CSF outflow and cause cerebral venous congestion (41, 42). Cortical crowding and compression at the top of the brain (resulting from the upward brain shift (16)) may impede CSF flow through the subarachnoid space along the superior cortical surface and/or impair CSF resorption into the superior sagittal sinus via the arachnoid granulations (19). Ventricular enlargement may thus reflect a compensatory response to accommodate increased intracranial CSF volume (2).

Also replicating our previous work (3, 16–18, 20), spaceflight resulted in widespread decreases in FW fractional volume around the vertex of the brain, primarily within the interhemispheric fissure and surrounding sulci. Post-flight FW fractional volume increased around the lower temporal and frontal lobes, primarily within the lateral fissure (Figure S3).

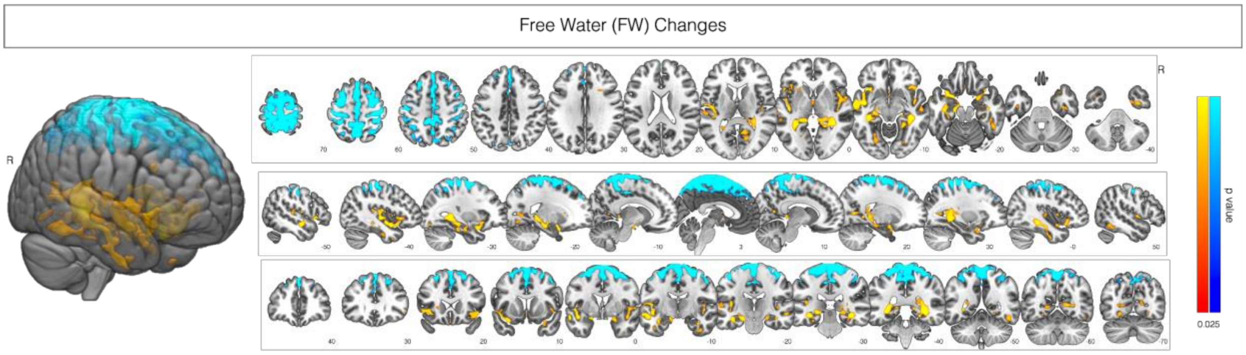


**Figure S3. Mean FW shifts following spaceflight.** Yellow clusters indicate regions where FW fractional volume increased from pre- to post-flight. Blue clusters indicate regions where FW fractional volume decreased from pre- to post-flight. FW fractional volume decreased around the vertex of the brain, primarily within the interhemispheric fissure and surrounding sulci. FW fractional volume increased around the lower temporal and frontal lobes, primarily within the lateral fissure. Clusters were FWE corrected at $p < 0.05$, two tailed. Clusters are overlaid on a MNI standard space template. R indicates the right hemisphere.
